# The different effect of pharmacological or low-doses of IFN-γ in endothelial cells are mediated by different intracellular signaling pathways

**DOI:** 10.1101/2022.09.13.507745

**Authors:** Alessandra Cazzaniga, Vincenzo Miranda, Sara Castiglioni, Jeanette A.M. Maier

**Affiliations:** Dipartimento di Scienze Biomediche e Cliniche L. Sacco, Università di Milano, 20157 Milan, Italy; Clinical Research Unit, GUNA S.p.a., Via Palmanova, 71, 20132 Milan, Italy

**Keywords:** interferon gamma, endothelial cells, ERK, STAT, migration

## Abstract

Interferon (IFN)-γ is a proinflammatory cytokine with a crucial role in intercellular communication during innate and acquired immune responses. IFN-γ interacts with many cell types, among which endothelial cells. Here, we show that pharmacological and low-dose kinetically activated (SKA) IFN-γ exert different effects on endothelial cells by activating different signal transduction pathways. Pharmacological concentrations of IFN-γ activate JAK/STAT pathway, inducing the overexpression of the CDKN1A p21, which in turn inhibits cell growth. Conversely, low-dose SKA IFN-γ does not activate the canonical JAK/STAT pathway but induces the phosphorylation of ERK. ERK activation is responsible for the induction of endothelial cell migration. Interestingly, ERK activation occurs only in the presence of kinetically activate low-dose IFN-γ, underlying the importance of mechanical forces to potentiate IFN-γ activity.

## 1. Introduction

The endothelium is a dynamic, disseminated organ which performs essential functions. In addition to acting as a physical barrier between the blood and the tissues, vascular endothelial cells (EC) modulate metabolic homeostasis, control coagulation and vascular tone. Moreover, EC play a central role in the regulation of local immune and inflammatory reactions. Vascular EC synthesize and release components of the complement cascade ^1^. Moreover, EC are a source and a target of chemokines and cytokines, and, when activated, express both Toll-like receptors (TLRs) and nucleotide-binding oligomerization domain (NOD)-like receptors (NLRs) as well as adhesion molecules for leukocytes ^2^, thus regulating immune cell recruitment to specific inflammatory sites. For this reason, EC were proposed as conditional innate immune cells ^2^. Because of their location, EC are one of the first targets of cytokines circulating in the blood stream. They respond to interferon (IFN)-γ, a proinflammatory cytokine crucial in intercellular communication during innate and acquired immune responses ^3^. Indeed, like most cell in the body, EC express the IFN-γ receptor (IFN-γR) ^3^. Its engagement by the cytokine activates the Janus Kinases (JAK)-1/2, followed by tyrosine phosphorylation of the cytoplasmic tail of the IFN-γR generating a docking site for Signal transducer and activator of transcription (STAT)-1 which is then phosphorylated by JAK2. Phosphorylated STAT-1 dimer translocates to the nucleus and binds to the γ-activated sequence elements (GAS elements) within the promoters of IFN-γ-responsive genes ^3^. In EC, IFN-γ modulates the expression of hundreds of genes ^4^, among which TNF-related apoptosis-inducing ligand (TRAIL), guanylate binding protein (GBP)1, and the chemokines CXCL10 and 11 ^4^, all involved in the inhibition of angiogenesis. This transcriptional program induced by IFN-γ might account for the widely reported inhibition of endothelial proliferation *in vitro* ^5–7^ as well as for its angiostatic effects *in vivo* ^8^. It has been shown that IFN-γ induces non apoptotic blood vessel regression in development, wound healing, remodeling of uterine vessels during pregnancy ^9–11^ as well as in experimental models of cancer ^8^. In these experimental settings the concentrations of IFN-γ utilized are much higher than the ones detected *in vivo* ^12–14^. Evidence is accumulating that ligand concentrations from 10^−18^ M to 10^−24^ M induce biological responses in various biological systems ^15^. However, the quantitative insight into the number of cytokine molecules needed to activate a target cell has been overlooked. A recent report demonstrates that interleukin (IL)-6 signaling requires only few IL-6 molecules ^16^. This is likely to be true for most cytokines, including IFN-γ, in physiological conditions. It should also be recalled that, in spite of its enormous therapeutic potential in immune diseases and in cancer, the use of IFN-γ is limited by its dose-dependent side effects ^17^. We have previously shown that very low-doses of IFN-γ exert an immunomodulatory action in Jurkat cells ^18^. Here we compare the effects of different concentrations of IFN-γ on the behavior of human EC, the first cells exposed to circulating IFN-γ.

## 2. Materials and Methods

### 2.1 Cell culture

HUVEC were from Lonza (Lonza, Basilea, Switzerland) and cultured in EBMTM Basal Medium (Lonza) supplemented with EGMTM Endothelial Cell Growth Medium SingleQuotsTM (Lonza) on 2% collagen-coated dishes. HUVEC were treated with pharmacological (10 ng/ml) or low-dose (10 pg/ml) concentrations of IFN-γ (PromoCell, Heidelberg, Germany) subjected or not to sequential kinetic activation (SKA). SKA IFN-γ was prepared by GUNA Laboratories (GUNA S.p.a., Milan, Italy) using a standardized method ^19^. IFN-γ was activated by sequential serial dilution (1:100) in 30% hydroalcoholic solution and kinetically energized by a shaking procedure (vertical shaking; 10 cm motion range; shaking speed corresponding to 100 oscillations in 10 seconds). Possible contaminations were excluded by HPLC (not shown). For proliferation experiment, 25×10^3^ cells/cm^2^ were seeded in 24-well plates and treated with pharmacological or low-dose concentrations of IFN-γ. After different time points, the cells were stained with Trypan blue solution (0.4%) (Sigma Aldrich), and the viable cells were counted using a cell counter. The experiments were performed three times in triplicate.

### 2.2 qRT-PCR

Total RNA was extracted by the PureLinkRNA MiniKit (Thermo Fisher Scientific). Single-stranded cDNA was synthesized from 1 μg RNA in a 20 μl final volume using the High-Capacity cDNA Reverse Transcription Kit with RNase inhibitor (Thermo Fisher Scientific), according to the manufacturer’s instructions. Real-time PCR (qRT-PCR) was performed three times in triplicate using the CFX96 Touch Real-Time PCR Detection System (Bio-Rad, Hercules, California, USA) instrument utilizing the TaqMan Gene Expression Assay (Life Technologies, Monza, Italy). The following primers (Thermo Fisher Scientific) were used: Hs00355782_m1 (cyclin-dependent kinase inhibitor 1 - *CDKN1A*) and Hs99999905_m1 (Glyceraldehyde-3-Phosphate Dehydrogenase - *GAPDH*) was used as an internal reference gene. Relative changes in gene expression were analyzed by the 2^-ΔΔCt^ method.

### 2.3 Western blot analysis

HUVEC were lysed in lysis buffer (50 mM Tris-HCl pH 8.0, 150 mM NaCl, 1 mM EDTA, 1% Nonidet P40 Substitute). Protein concentration was determined using the Bradford reagent (Sigma Aldrich, St. Louis, MO, USA). Equal amounts of proteins were separated by SDS-PAGE and transferred to nitrocellulose membranes by using Trans-Blot Turbo™ Transfer Pack (Bio-Rad, Hercules, CA, USA). Western blot analysis was performed using primary antibodies against p-ERK (Cell Signaling Technologies Danvers, MA, USA), ERK1/2 and GAPDH (Santa Cruz Biotechnology, Dallas, TX, USA). Secondary antibodies conjugated with horseradish peroxidase (Amersham Pharmacia Biotech Italia, Cologno Monzese, Italy) were used. The immunoreactive proteins were detected with Clarity™ Western ECL substrate (Bio-Rad) and images were captured with a ChemiDoc MP Imaging System (Bio-Rad). Densitometry of the bands was performed with the software ImageJ (National Institute of Health, Bethesda, MD, USA). The Western blots shown are representative and the densitometric analysis was performed on three independent experiments ± standard deviation (SD)

### 2.4 ELISA

The PathScan PHOSPHO-STAT-1 (Tyr701) Sandwich ELISA Kit (Cell Signaling, Pero-MI, Italy; Catalog Number: 7234) was used to measure the activation of STAT-1 according to the manufacturer’s instructions. InstantOne ELISA ERK1/2 (Total/Phospho) (Invitrogen, Thermo Fisher Scientific; Catalog Number 85-86013-11) was used to analyze ERK1/2 phosphorylation according to the manufacturer’s instructions. All ELISAs were performed three times, and each sample was measured in triplicate.

### 2.5 Migration assay

Cell migration was determined using an *in vitro* model of wound repair as previously described ^20^. HUVEC were grown in 24-well plates to confluence. After 16 h of starvation in medium without FBS and Endothelial Cell Growth Factor, the monolayer was wounded and treated with pharmacological or low-dose concentrations of IFN-γ. 10 ng/ml of vascular endothelial growth factor (VEGF) (PeproTech London UK) were used as positive control. The cells were stained with crystal violet to visualize the width of the wound and images were captured with a 4X objective using a phase contrast microscope ^18^. The wound area was calculated by ImageJ software and expressed using an arbitrary value scale. The cells were also fixed in phosphate buffered saline containing 4% paraformaldehyde and 2% sucrose (pH 7.6) and stained with rhodamine-labeled phalloidin and DAPI (4′,6-diamidino-2-phenylindole) to visualize the cytoskeleton and the nuclei, respectively. The images were acquired using 20X objective by FLoid Cell Imaging Station (Thermo Fisher Scientific). In some experiments, after starvation, HUVEC were treated with 30 μM ERK inhibitor III (Calbiochem, San Diego, CA, USA) 1 h before performing the wound and the treatment with IFN-γ. To obtain a transient downregulation of ERK2 in HUVEC, we used Lipofectamine RNAiMAX (Thermo Fisher Scientific, Waltham, MA, USA) according to the manufacturer’s recommendations in combination with the stealth siRNAs for *ERK2* (FlexiTube GeneSolution GS5594 for MAPK1, Qiagen, Hilden, Germany). Non-silencing, scrambled sequences were used as controls. After transfection, the cells were starved for 16 h and then wounded and treated with IFN-γ. At the end of the experiment, wound area was calculated by ImageJ software on crystal violet - stained cells and expressed using an arbitrary value scale. The experiments were performed three times in triplicate. Data are shown as the mean ± SD.

### 2.6 Statistical analysis

Data are expressed as the mean ± SD. The data were analyzed using one-way ANOVA. The *p*-values deriving from multiple pairwise comparisons were corrected using the Bonferroni method. Statistical significance was defined as *p*-value ≤ 0.05. In the figures, ** *p* ≤ 0.01; *** *p* ≤ 0.001.

## 3. Results

### 3.1 Low-dose IFN-γ does not affect HUVEC proliferation

Several studies demonstrate that IFN-γ inhibits endothelial growth ^5–7^. We compared the proliferative response of HUVEC exposed either to pharmacological (10 ng/ml) or low (10 pg/ml) concentrations of IFN-γ, subjected or not to sequential kinetic activation (SKA). Fig. 1A shows that 10 ng/ml IFN-γ, both SKA or not, inhibited HUVEC proliferation. Growth inhibition is associated with the overexpression of the cyclin-dependent kinase inhibitor 1 (*CDKN1A*), which codes for p21, as detected by qRT-PCR (Fig. 1B). Conversely, 10 pg/ml IFN-γ, SKA or not, had no effects on HUVEC proliferation and *CDKN1A* expression (Fig. 1A-B).

**Fig. 1.**
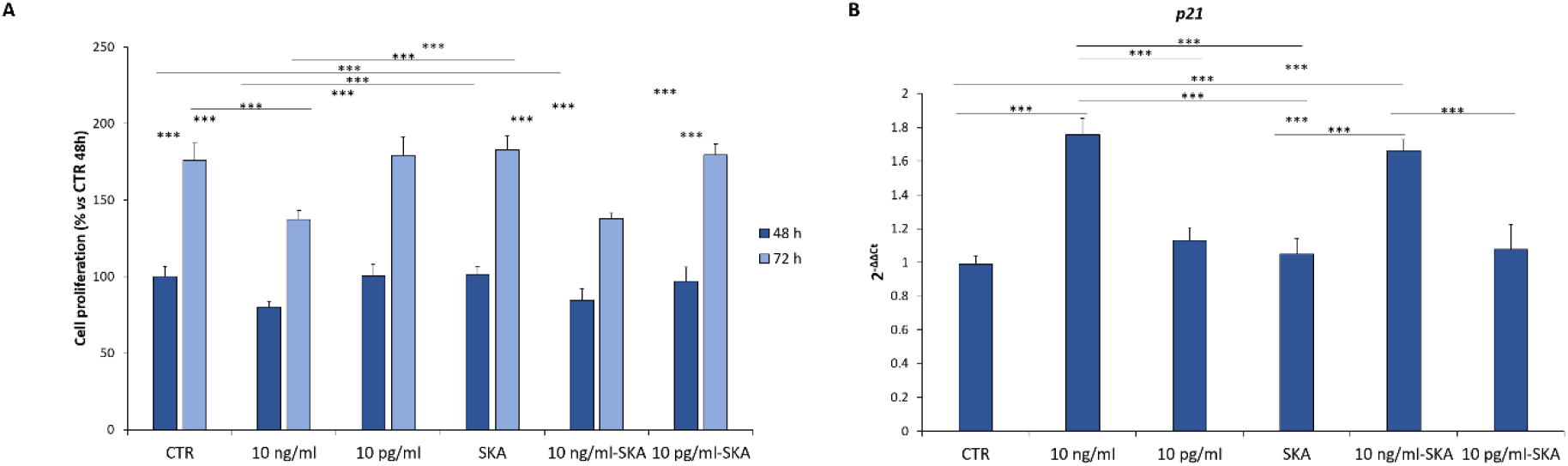
Pharmacological but not low concentrations of IFN-γ inhibit HUVEC proliferation. HUVEC were treated with pharmacological (10 ng/ml) or low (10 pg/ml) concentrations of IFN-γ, subjected or not to sequential kinetic activation (SKA). CTR is the untreated control. (A) The cells were counted after 48 and 72 h. The results are the mean of four separate experiments performed in triplicate. (B) After 16 h of treatment, qRT-PCR was performed on RNA extracted from the cells as described in the methods. The experiments were performed three times in triplicate ± SD.

### 3.2 Low-dose SKA IFN-γ increases HUVEC migration

Endothelial migration is an important event in angiogenesis ^21^ and in the repair of damaged endothelial lining caused by toxic injury or harmful mechanical stimuli ^22^. Therefore, we used wound assay to analyze the migratory capacity of HUVEC in the presence of pharmacological or low-dose concentrations of IFN-γ subjected or not to SKA. 10 ng/ml of vascular endothelial growth factor (VEGF) were used as positive control. Fig. 2A shows photos obtained in a representative experiment after staining the cells with crystal violet. The cells were also stained with rhodamine-labeled phalloidin to detect the cytoskeleton and DAPI to visualize the nuclei. 10 pg/ml SKA IFN-γ and VEGF induce polarized cell elongation (highlighted by the white arrows and magnified in the lower box), which is typical of endothelial migratory phenotype ^20^. Quantification of the wound width is reported in Fig. 2B, which shows that 10 pg/ml SKA IFN-γ increased HUVEC migration as much as VEGF. Interestingly, HUVEC migration was not modulated by non SKA low-dose IFN-γ or pharmacological concentrations of IFN-γ.

**Fig. 2.**
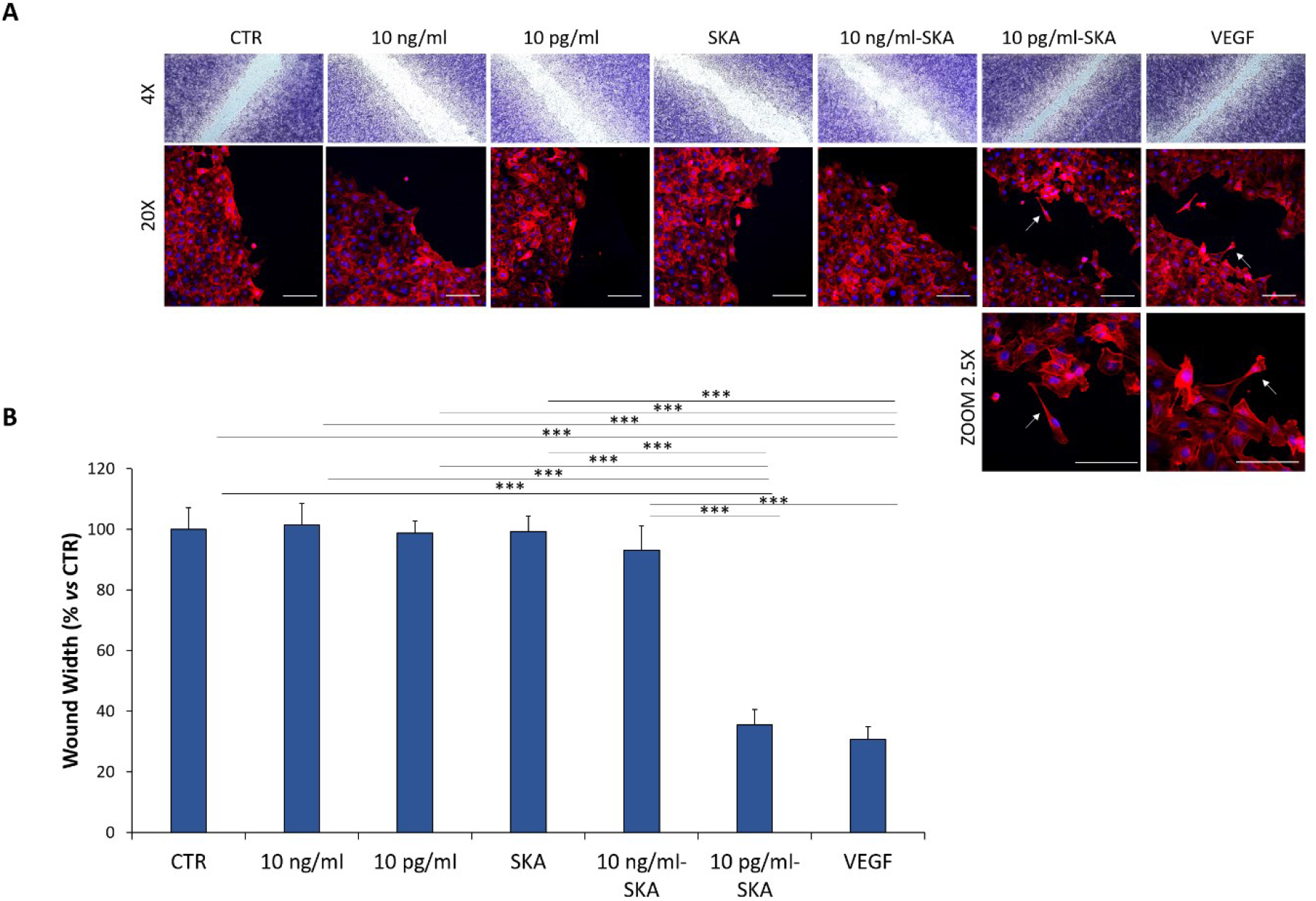
Low-dose SKA IFN-γ induces HUVEC migration. HUVEC were grown in 24-well plates to confluence. After 16 h of starvation, the monolayer was wounded and the cells were treated with pharmacological (10 ng/ml) or low (10 pg/ml) concentrations of IFN-γ, subjected or not to sequential kinetic activation (SKA). CTR is the untreated control. VEGF (10 ng/ml) was used as a positive control. (A) After 24 h, the cells were stained with crystal violet and images acquired with a 4X objective using a phase contrast microscope. Cytoskeleton was visualized after staining with fluorescein-labeled phalloidin and nuclei with DAPI. Scale bar: 100 μm. A representative experiment is shown. The white arrows indicate the cells with a migratory phenotype which are magnified in the lower box. (B) The wound area was calculated by ImageJ software. Data are shown as the mean of three experiments ± SD.

### 3.3 Low-dose SKA IFN-γ activates ERK

We then investigated the JAK/STAT pathway, which is the main intracellular signaling transducer in IFN-γ-treated cells. We cultured HUVEC in the presence of pharmacological or low-dose concentrations of IFN-γ, subjected or not to SKA, and analyzed STAT-1 phosphorylation by ELISA. Fig. 3A shows that 30 min of treatment with 10 ng/ml IFN-γ both SKA or not SKA is sufficient to induce STAT-1 phosphorylation. Conversely, no phosphorylation of STAT-1 was observed after treating HUVEC with low-dose IFN-γ. Since it is known that other pathways, including MAP kinase, co-operate with or act in parallel to JAK/STAT pathway to transduce IFN-γ signal ^23^, we analyzed whether MAP kinase signaling is involved. Both western blot and ELISA demonstrate that 10 pg/ml-SKA IFN-γ induced ERK phosphorylation (Fig. 3B). ERK was not phosphorylated by non SKA low-dose and pharmacological concentration of IFN-γ.

**Fig. 3.**
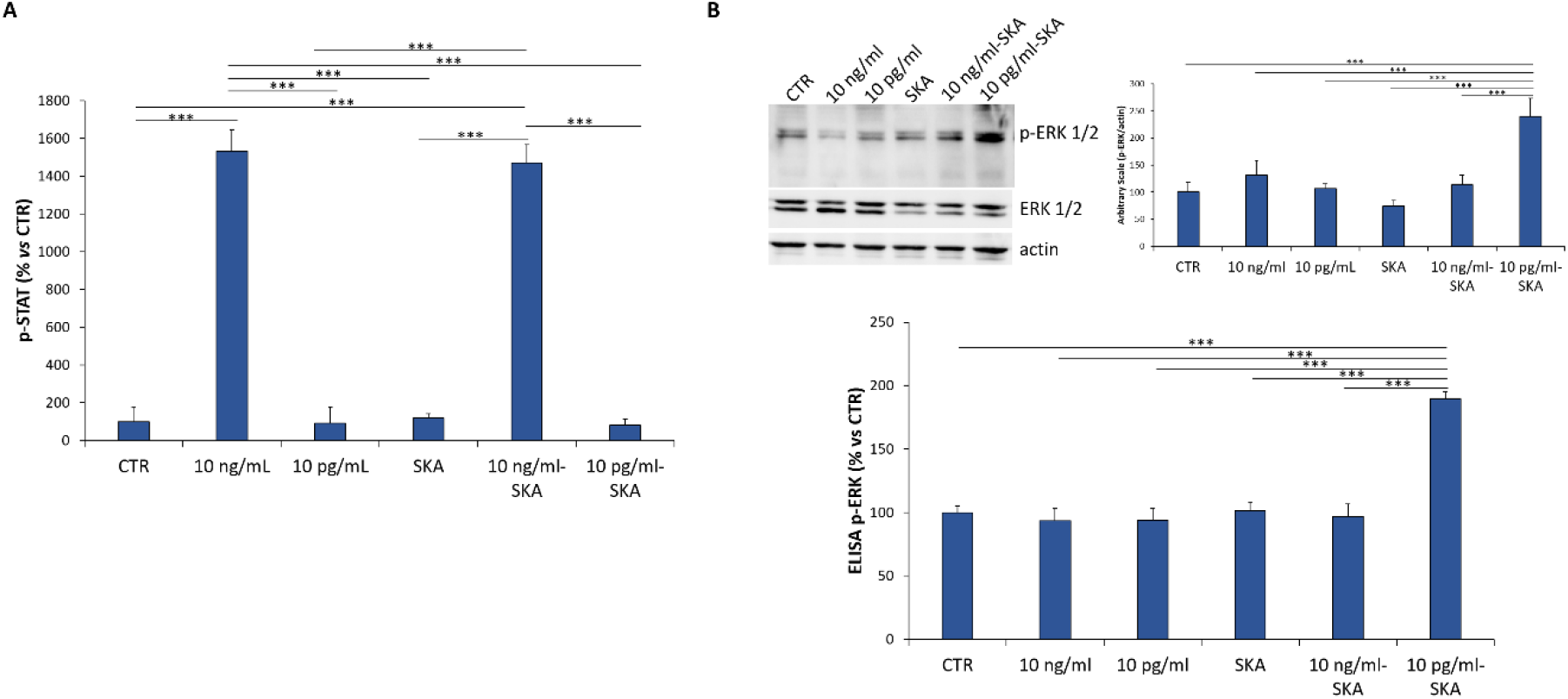
Low-dose SKA IFN-γ activates MAP kinase pathway. (A) The phosphorylation of STAT-1 (p-STAT) was evaluated by ELISA on HUVEC after 30 min of treatment with pharmacological (10 ng/ml) or very low (10 pg/ml) concentrations of IFN-γ, subjected or not to sequential kinetic activation (SKA). (B) After 16 h of starvation, HUVEC were treated for 30 min with the different concentrations of IFN-γ. Then the cells were lysed and the phosphorylation of ERK1/2 was analyzed by western blot (upper panel) and ELISA (lower panel). Actin was used as a control of loading. A representative blot and densitometry obtained by ImageJ on three independent experiments ± SD are shown. The ELISA results are the mean of three experiments in triplicates ± SD. CTR is the untreated sample.

### 3.4 ERK phosphorylation mediates the increase of endothelial migration after treatment with low-dose SKA IFN-γ

To understand if ERK activation mediates low-dose SKA IFN-γ-induced HUVEC migration, we inhibited ERK activation using a specific pharmacological inhibitor, i.e. ERK inhibitor III. We then analyzed the migration of HUVEC cells in the presence of pharmacological or low-dose concentrations of IFN-γ. ERK inhibition prevents the increase of HUVEC migration induced by low-dose SKA IFN-γ (Fig. 4). To reinforce these data, we silenced *ERK* using specific siRNA. Fig 4 shows that siRNA significantly inhibited low-dose SKA IFN-γ-induced migration (Fig. 4).

**Fig. 4.**
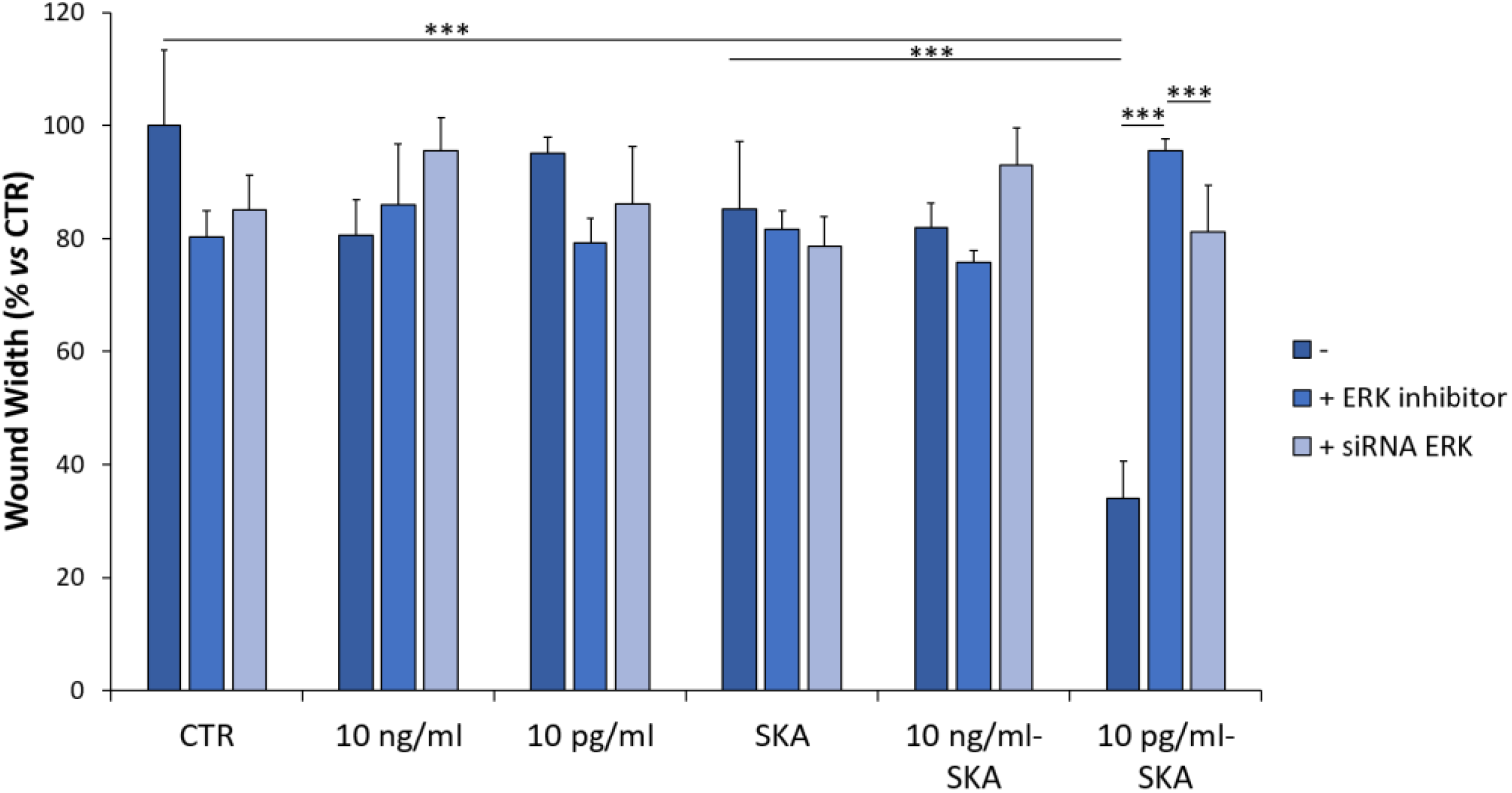
ERK inhibition blocks HUVEC migration induced by low-dose SKA IFN-γ. HUVEC were grown in 24-well plates to confluence and wound assay was performed in HUVEC treated with the ERK inhibitor III (30 μM) or silenced for ERK as described in the methods. The experiments were performed in triplicate. Data are shown as the mean ± SD. CTR is the not treated control sample.

## 4. Discussion

The use of low-dose kinetically activated cytokines is taking its first steps in *in vitro* studies as well as in clinical settings ^24^. *In vivo*, the principal aim is the harmonization of cellular function to maintain or restore homeostasis. Different kinetically activated cytokines have yielded promising results in patients with psoriasis ^25^, atopic dermatitis ^26^ and rheumatoid arthritis ^27^. Concerning IFN-γ, low-doses of SKA IFN-γ potentiated the activity of natural killer cells isolated from patients with early-stage colon cancer ^28^. In Jurkat cells, low-dose SKA IFN-γ exerted an immunomodulatory action by tuning signal transduction and gene expression ^18^.

Here we compare the effects of pharmacological and low-dose IFN-γ, kinetically activated or not, in cultured EC. We show that pharmacological and low-dose SKA IFN-γ exert different effects in HUVEC by activating different signal transduction pathways (Fig. 5). 10 ng/ml IFN-γ, subjected or not to kinetic activation, inhibited cell growth by upregulating the CDKN1A p21, in agreement with previous results obtained in HUVEC ^29^ and other cell types ^30^. It is known that activated STAT-1 specifically binds the conserved STAT-responsive elements in the promoter of the gene encoding p21, resulting in the overexpression of its messenger RNA. Consequently, it is STAT-1 activation that mediates IFN-γ-dependent growth inhibition, as demonstrated in HUVEC exposed to siRNA targeting *STAT-1* ^29^. Consistently, kinetically activated or not 10 pg/ml of IFN-γ, which do not activate STAT-1, do not influence cell proliferation nor the expression of CDKN1A. Intriguingly, only SKA low-dose IFN-γ stimulates cell migration through the activation of ERK. Indeed, siRNA targeting *ERK* as well as the ERK inhibitor III prevent the stimulation of cell migration induced by low-dose SKA IFN-γ. It is noteworthy that ERK is central in endothelial migration as shown in primary endothelial cells derived from ERK knock out mice ^31^. Two main questions arise at this point.

**Fig. 5.**
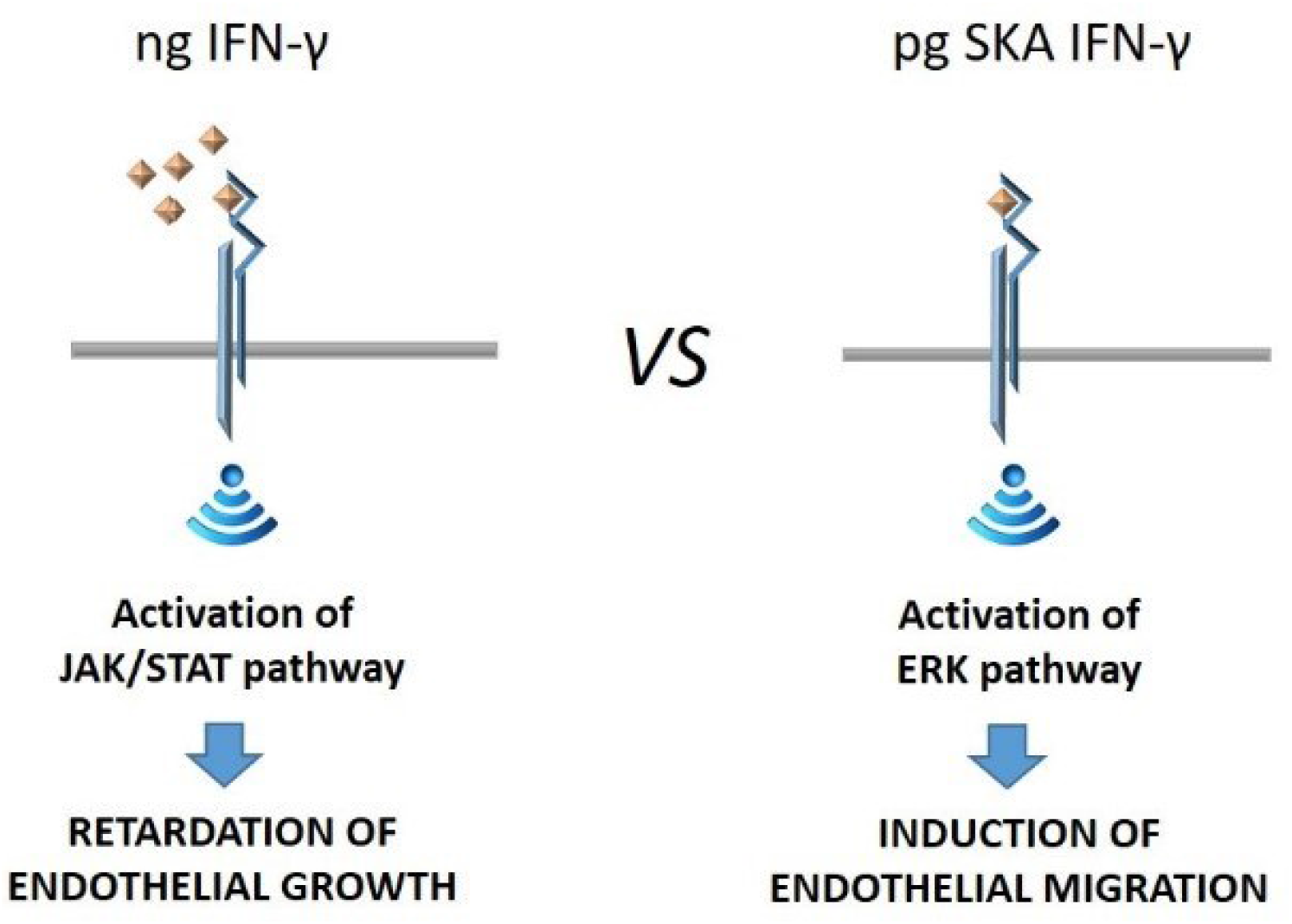
The mechanisms of action of pharmacological vs low-dose SKA IFN-γ. The cartoon summarizes the findings of our work.

The first is about the involvement of ERK in the signal cascade, and the second one concerns the need to kinetically activate the low-dose cytokine to potentiate its action. While STAT-1 is essential in cell response to IFN-γ, several studies have also demonstrated the existence of STAT-1 independent pathways, often in a cell type specific fashion ^32^. As examples, we recall that ERK is rapidly activated in IFN-γ treated neurons ^32^ and ERK-dependent STAT-1 phosphorylation maximizes STAT-1 activity in macrophages ^33^. Therefore, it is known that ERK contributes to the signaling activated by pharmacological concentrations of IFN-γ. The novelty of our results lies in the evidence that low-dose SKA IFN-γ does not activate the canonical STAT-1 pathway while it phosphorylates ERK, which mediates the stimulation of endothelial migration. We hypothesize that low-doses of SKA IFN-γ unmask the intervention of ERK pathway, which is otherwise overwhelmed by STAT-1 signaling. Therefore, the signal transduction pathway operated by ERK might be more sensitive to the action of minimal doses of the cytokine.

Since pharmacological concentrations of IFN-γ not only inhibit growth but also stimulate apoptosis of HUVEC ^34^, while low concentrations maintain normal proliferation rate and increase migration, we propose that these apparently conflicting results might be framed in the perspective of the hormetic dose responses, which means that low concentrations of potentially harmful molecules exert beneficial effects ^35^. Hormesis is now envisioned as an adaptive cellular response which contributes to the maintenance of homeostasis by enhancing biologic plasticity ^36^. Since the exceptional sensitivity to low concentrations of ligands is maintained from unicellular to mammalian organisms, hormesis might have played an important role in evolution as a survival strategy ^15^. Evidence is accumulating about the effects of low-doses of biological active compounds ^15^. A recent example comes from studies on relaxin, which has been shown to be active in a wide range of doses, from attomolar to millimolar concentrations ^37^. This seems to be due to the fact that the relaxin receptor, a G-protein coupled receptor, preassembles into a large signaling complex that facilitates its activation by very low concentrations of relaxin ^37^. On these bases, we speculate that low-dose IFN-γ might stimulate ERK phosphorylation by binding to particular domains of the plasma membrane enriched in IFN-γ receptors.

It should be recalled that most of the experimental studies are performed using concentrations much higher than the ones detected *in vivo* ^12–14^. The quantitative insight into the number of cytokine molecules needed to activate a target cell has not attracted attention until recently. In 2020 a report demonstrated that IL-6 signaling requires only few IL-6 molecules ^16^. This is likely to be true for most cytokines, including IFN-γ, in physiological conditions.

However, in our experimental model only SKA low-dose IFN-γ activates ERK and stimulates HUVEC migration. A recent study has shown that mechanical exposure alters the physical-chemical characteristics of water with consequences on the infrared emission spectra of ultra-high diluted IFN-γ solutions ^38^. On these bases, we hypothesize that mechanical forces might determine conformational changes of IFN-γ itself or its receptor to optimize their binding and maximize cell response.

In conclusion, in HUVEC STAT-1 serves as the principal intracellular signal activated by nanograms of IFN-γ, independently of mechanical stimulation, whereas ERK mediates the increase of endothelial migration in response to low-dose kinetically activated IFN-γ (Fig. 5).

## Acknowledgements

We thank Roberta Scrimieri for her help in some experiments.

## Author Contributions

Conceptualization, Vincenzo Miranda, Sara Castiglioni and Jeanette A.M. Maier; Formal analysis, Alessandra Cazzaniga; Funding acquisition, Jeanette A.M. Maier; Investigation, Alessandra Cazzaniga; Methodology, Alessandra Cazzaniga; Supervision, Sara Castiglioni; Validation, Alessandra Cazzaniga, Vincenzo Miranda, Sara Castiglioni and Jeanette A.M. Maier; Writing – original draft, Jeanette A.M. Maier; Writing – review & editing, Sara Castiglioni and Jeanette A.M. Maier. All authors have read and agreed to the published version of the manuscript.

